# Small CD4 mimetics sensitize HIV-1-infected macrophages to antibody-dependent cellular cytotoxicity

**DOI:** 10.1101/2022.06.30.498265

**Authors:** Annemarie Laumaea, Lorie Marchitto, Guillaume Beaudoin-Bussières, Shilei Ding, Romain Gasser, Debashree Chatterjee, Gabrielle Gendron-Lepage, Halima Medjahed, Hung-Ching Chen, Amos B Smith, Haitao Ding, John C. Kappes, Beatrice H. Hahn, Frank Kirchhoff, Jonathan Richard, Ralf Duerr, Andrés Finzi

**Affiliations:** Centre de Recherche du CHUM, QC H2X 0A9, Canada; Département de Microbiologie, Infectiologie et Immunologie, Université de Montréal, Montreal, QC H2X 0A9, Canada; Department of Chemistry, School of Arts and Sciences, University of Pennsylvania, Philadelphia, PA 19104-6323, USA; Department of Medicine, University of Alabama at Birmingham, Birmingham, AL, USA; Birmingham Veterans Affairs Medical Center, Research Service, Birmingham, AL 35233, USA; Departments of Medicine and Microbiology, Perelman School of Medicine, University of Pennsylvania, Philadelphia, PA 19104-6076, USA; Institute of Molecular Virology, Ulm University Medical Center, Ulm, Germany; Department of Microbiology, New York University School of Medicine, New York, NY 10016, USA

**Keywords:** CD4, BST-2, HIV Envelope, Accessory proteins, Open conformation, Closed conformation, bNAbs, nnAbs, Antibody Dependent Cellular Cytotoxicity

## Abstract

HIV-1 Envelope (Env) conformation determines the susceptibility of infected CD4^+^ T cells to Antibody Dependent Cellular Cytotoxicity (ADCC). Upon interaction with CD4, Env adopts more “open” conformations, exposing ADCC epitopes. HIV-1 limits Env-CD4 interaction and protects infected cells against ADCC by downregulating CD4 via Nef, Vpu and Env. Limited data exists however of the role of these proteins in downmodulating CD4 on infected macrophages and how this impacts Env conformation. While Nef, Vpu and Env are all required to efficiently downregulate CD4 on infected CD4^+^T cells, we show here that any one of these proteins is sufficient to downmodulate most CD4 from the surface of infected macrophages. Consistent with this finding, Nef and Vpu have a lesser impact on Env conformation and ADCC sensitivity in infected macrophages compared to CD4^+^T cells. However, treatment of infected macrophages with small CD4-mimetics expose vulnerable CD4-induced Env epitopes and sensitize them to ADCC.

## Introduction

Being the only viral protein exposed on the surface of infected cells, the HIV-1 Envelope glycoprotein (Env) is the main antigen targeted by neutralizing and non-neutralizing antibodies. Indeed, broadly neutralizing antibodies (bNAbs) have been shown to efficiently mediate Antibody Dependent Cellular Cytotoxicity (ADCC) against productively infected cells (Anand et al., 2021; Bruel et al., 2016; Richard et al., 2018; von Bredow et al., 2016). Most of these studies largely focused on ADCC responses against infected CD4^+^T cells. However, comparatively little is known about the conformation of Env at the surface of infected macrophages and consequently its impact on responses against this cell type, with one recent study reporting resistance of infected macrophages to NK-mediated killing using select bNAbs (Clayton et al., 2021).

Macrophages which are distributed throughout the body are highly heterogeneous (Reviewed in (Qian et al., 2019)). Tissue location of macrophage populations (blood, brain, gut, lungs, liver) is intimately associated with their ontogeny and several lineages/subsets have been described based on their developmental origins; Embryonic Yolk Sac and Fetal Liver precursors vs Bone marrow progenitors (Reviewed in (Ginhoux and Guilliams, 2016)). To date the relevance of HIV infection in the different subsets is poorly defined, though evidence support microglia, a tissue resident subset, as an important target for HIV in the brain (Borrajo et al., 2021; Cenker et al., 2017; Wallet et al., 2019). Despite the susceptibility of macrophages to HIV infection (Gendelman et al., 1988; Ho et al., 1986; Jambo et al., 2014; Nicholson et al., 1986) there exists important differences between CD4^+^T cells and macrophages that could influence the efficiency of eliminating these cells. An important difference is cell surface levels of CD4 (Lee et al., 1999) which have been shown to be important in the exposure of ADCC epitopes on CD4^+^T cells (Prévost et al., 2022; Prevost et al., 2018; Veillette et al., 2014). Here we used monocyte-derived macrophages (MDM) as a model to better understand the role of HIV-1 Nef, Vpu and Env on CD4 downregulation in this cell type and studied how this impacts Env conformation and susceptibility to ADCC.

## Results

### HIV-1 mediated CD4 downregulation in macrophages

HIV-1 uses Vpu, Env and Nef to downregulate CD4 from the surface of primary CD4^+^ T cells (Benson et al., 1993; Chen et al., 1996; Dalgleish et al., 1984; Delwart and Panganiban, 1989; Klatzmann et al., 1984; Wildum et al., 2006). To evaluate whether this was also the case for macrophages, which express comparatively little CD4 (Lee et al., 1999), we infected monocyte-derived macrophages and autologous CD4^+^T cells with the wild-type (WT) full-length infectious molecular clone (IMC) of the macrophage tropic, R5 isolate, HIV-1_AD8_. The relative contribution towards CD4 downregulation by these viral proteins was measured by infecting cells with mutant IMCs unable to express Nef (N-), Vpu (U-) or both accessory proteins (N-U-). Additionally, the impact of Env on CD4 downregulation was measured with IMCs expressing an Env variant containing a mutation in the CD4 binding site (CD4BS) that prevents Env interaction with CD4 (Kwong et al., 1998; Veillette et al., 2015) (D368R; N^-^U^-^D368R). In both cell types, the maximal CD4 downregulation was achieved with the WT IMC (Figure 1A; Table S1). Infection with N-, U- and N-U- but not D368R significantly disrupted CD4 downregulation in primary CD4^+^T cells. Of note, Env contribution to CD4 downregulation was evident when the D368R mutation was added to the N-U- IMC (N-U- D368R). In agreement with previous observations (Veillette et al., 2015; Veillette et al., 2014), only in this context infected cells expressed the same levels of CD4 than uninfected cells (Figure 1A and B). Neither the abrogation of Nef and/or Vpu nor the CD4BS mutation completely restored CD4 surface levels to those observed in uninfected cells. In both cell types, all three mechanisms of CD4 downregulation (i.e., Nef, Vpu and Env, N-U-D368R) needed to be disrupted in order to present comparable levels of CD4 expression than mock infected cells (Figure 1A and B). As expected, in CD4^+^ T cells deletion of Nef or Vpu resulted in a significant increase of cell surface levels of CD4 (Figure 1A and B). The same deletions didn’t impact CD4 levels at the surface of infected macrophages, which remained the same than in wt-infected cells. Both Nef and Vpu had to be deleted (N-U-) in order to observe a significant increase in cell surface levels of CD4 in macrophages (Figure 1A and B). The lower overall CD4 expression on macrophages, as previously reported (Lee et al., 1999) and supported herein (Figure 1A), likely explains this phenotype. To ensure that this phenotype was not restricted to HIV-1_AD8_, we performed the analogous experiments with two additional R5 (HIV-1_JRFL_, HIV-1_YU2_) and one dual tropic (HIV- 1_CH77_) IMCs. Similar phenotypes were observed (Figure 1B; Figure S1-E-G; Table S1).

**Figure 1.**
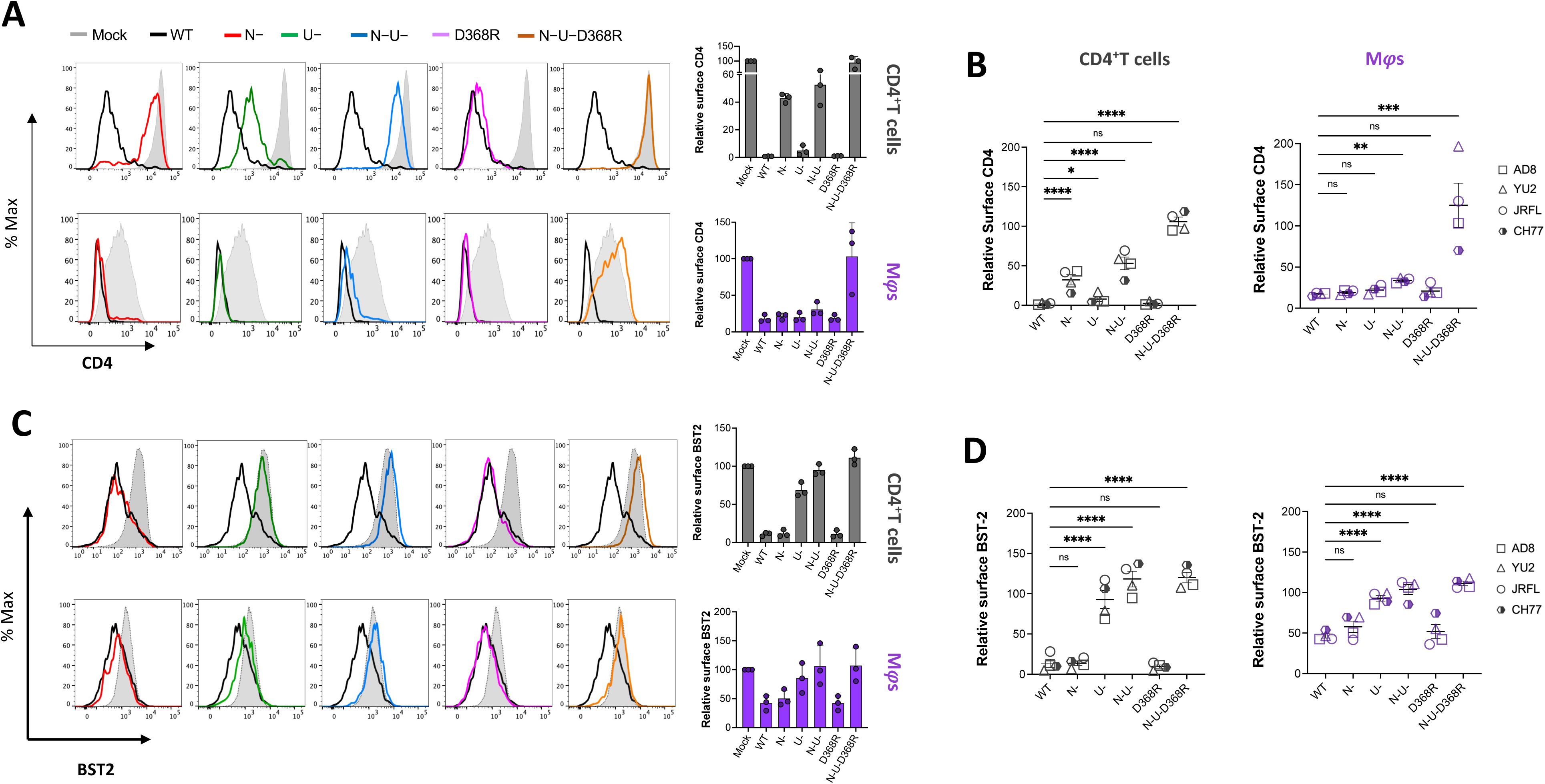
CD4 and BST2 downregulation in HIV-1-infected primary CD4^+^ T cells and autologous macrophages. Autologous CD4^+^T cells and macrophages were infected with the HIV-1_AD8_ panel of viruses [wt, nef-defective (N-), Vpu-defective (U-), or both Nef and Vpu (N-U-) and CD4BS mutant (D368R) and N-U-D368R mutant] and CD4 and BST2 levels were determined 48h later (CD4^+^T cells) or 5 days post infection (Macrophages). (**A**) Relative surface levels of CD4 on CD4^+^T cells and macrophages. (*Left*) Histograms depict representative staining of infected cells and bar graphs (*Right)* show percent fold change in CD4 expression relative to Mock (p24+/uninfected cells). (**B**) Summary of relative surface levels of CD4 on CD4^+^T cells and macrophages infected with a panel of viruses [wt, nef-defective (N-), Vpu-defective (U-), or both Nef and Vpu (N-U-) and CD4BS mutant (D368R) and N-U-D368R mutant] from HIV-1_AD8_, HIV-1_JR-FL_, HIV-1_CH77_ and HIV-1_YU2_. (**C**) Relative surface expression of BST2 on CD4^+^ T cells and macrophages. (*Left*) Histograms depict representative staining of infected cells and bar graphs (*Right)* show percent fold change in BST2 expression of p24^+^ relative to p24^-^ cells (p24+/p24-). (**D**) Summary of relative surface expression of BST2 on CD4^+^T cells and macrophages infected with a panel of viruses [wt, nef-defective (N-), Vpu-defective (U-), or both Nef and Vpu (N-U-) and CD4BS mutant (D368R) and N-U-D368R mutant] from HIV-1_AD8_, HIV-1_JR-FL_, HIV-1_CH77_ and HIV-1_YU2_. Statistical significance was tested using Ordinary One-way Anova (*p<0.05; **p<0.001; ***p<0.0001; ns, non-significant).

In HIV-1-infected CD4^+^ T cells, Nef and Vpu restrict ADCC responses by limiting Env-CD4 complexes that otherwise expose vulnerable epitopes recognized by antibodies commonly present in plasma from infected individuals (Alsahafi et al., 2015; Ding et al., 2016b; Veillette et al., 2015; Veillette et al., 2014). In addition to downregulating CD4, Vpu contributes to protection of infected cells from ADCC by downregulating the restriction factor BST-2, which traps viral particles and results in antigen accumulation at the cell surface (Alvarez et al., 2014; Arias et al., 2014; Veillette et al., 2015; Veillette et al., 2014). We therefore measured cell surface levels of BST-2 in infected CD4^+^T cells and autologous macrophages. In both cell types, Vpu was the principal viral determinant modulating BST-2 expression (Figure 1C). Similar results were obtained with the other IMCs tested (Figure 1D, Figures S1 E-G and Table S1). Thus, with respect to BST-2 modulation, Vpu performs parallel roles on CD4^+^T cells and macrophages. Of note, we observed that for HIV-1 AD8 and YU2, full recovery of BST-2 levels at the cell surface required Nef deletion (N-U-), in agreement with a recent report suggesting that Nef from some HIV-1 strains, including AD8 can downregulate BST-2 (Giese et al., 2020).

### Env conformation at the surface of infected primary CD4^+^ T cells and autologous macrophages

Using a panel of anti-Env antibodies (Table S2) and IMC variants (WT, N-, U-, N-U-, D368R and N-U-D368R), we evaluated Env conformation at the surface of infected macrophages and autologous CD4^+^T cells. In both cell types, recognition of infected cells with bNAbs having a conformational preference for the “closed” trimer (PG9, PGT121, 10.1074, 3BNC.117, PGT151) was modulated by Vpu. Vpu deletion (U-) alone or in combination with Nef (N-U-) enhanced recognition, in a manner that reflected overall amount of Env at the surface of infected cells, as measured with the conformational independent 2G12 antibody (Figure 2A). This is consistent with previous studies demonstrating that Vpu-mediated BST-2 downregulation prevents virion accumulation on the cell surface thereby reducing the overall levels of detectable Env (Alvarez et al., 2014; Arias et al., 2014; Richard et al., 2017; Veillette et al., 2015; Veillette et al., 2014). The impact of Vpu on cell recognition by the different bNAbs was markedly lower on infected macrophages compared to CD4^+^T cells (Figure 2).

**Figure 2.**
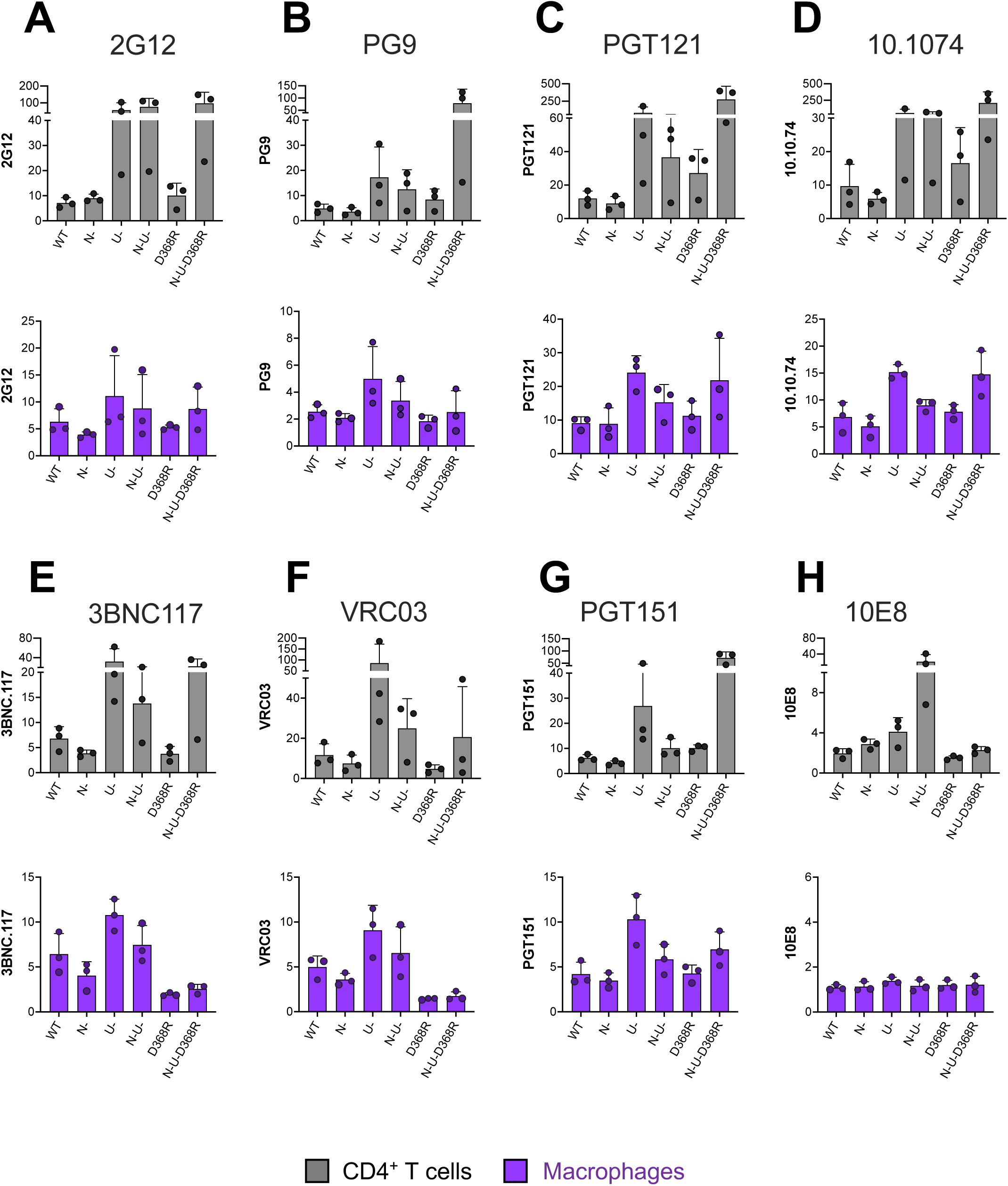
Env recognition at the surface of HIV-1-infected primary CD4^+^ T cells and autologous macrophages by a panel of bNAbs. Staining of CD4^+^T cells and macrophages infected with HIV-1_AD8_ panel of viruses [wt, nef-defective (N-), Vpu- defective (U-), or both Nef and Vpu (N-U-) and CD4BS mutant (D368R) and N-U- D368R mutant] with (**A**) 2G12 a conformation independent antibody; (**B**) the V1V2- apex antibody PG9; V3glycan antibodies (**C**) PGT121 and (**D**) 10.1074; CD4BS antibodies (**E**) 3BNC117 and (**F**) VRC03; (**G**) PGT151 which targets the gp120-gp41 interface; and the (**H**) anti-MPER antibody 10E8. Values represent fold change in MFI relative to mock.

PG9 recognizes V1V2 epitopes on the trimer apex (Doores and Burton, 2010; McLellan et al., 2011; Walker et al., 2009) and preferentially interacts with the “closed” Env. Indeed, interaction of Env with membrane-bound CD4 was shown to decrease PG9 interaction (Veillette et al., 2014). Supporting the existence of a “closed” conformation of the unliganded trimer, we observed similar levels of Env recognition by PG9 (Figure 2B) at the surface of WT-infected CD4^+^T cells and macrophages (Figure 2B). Interestingly, while CD4–Env disruption in the context of the N-U-D368R infected cells significantly enhanced binding of PG9 on CD4^+^T cells, this was not observed for macrophages (Figure 2B, *lower panel*). This phenotype was recapitulated with the HIV-1_CH77_ and HIV-1_YU2_ N-U-D368R mutants (Figure S2-B).

Recent work showed comparable staining of Env with the 3BNC117 CD4BS Ab on both cell types (Clayton et al., 2021) while others such as N6 (Clayton et al., 2021) and NIH-45-46 (Kek et al., 2021) showed better recognition of macrophages in comparison to CD4^+^T cells. Consistent with Clayton *et al,* (Clayton et al., 2021) we observed that 3BNC117 and VRC03 recognize Env similarly on both WT-infected CD4^+^T cells and macrophages (Figure 2E & F). These CD4BS antibodies displayed improved recognition of cells infected with a Vpu-virus (Figure 2E-F). This recognition was diminished by deleting Nef. In the absence of Nef there is more CD4 on the cell surface interacting with Env (Veillette et al., 2014), therefore occluding the CD4BS. The D368R variant slightly decreased recognition by these antibodies, as expected due to the role played by D368 for their interaction (Klein et al., 2013; Zhou et al., 2010). While disruption of the Env-CD4 interaction in the absence of Nef and Vpu (N-U- D368R) enhanced 3BNC117 recognition of infected CD4^+^T cells, this didn’t happen in macrophages. This is likely due to low amounts of CD4 on the surface of macrophages which might not be sufficient to compete with this CD4BS bNAb (Figure 2 E and F). When PGT151, an antibody that recognizes the interface between gp120 and gp41 (Falkowska et al., 2014) was tested, we observed efficient recognition of cells infected with a virus lacking Vpu or expressing Env D368R, but not when both Vpu and Nef were absent (Figure 2G). At the surface of infected cells, CD4 and Env are interacting on the same membrane and the CD4 domains D3-D4 may prevent PGT151 access of its epitope on Env, which is located beneath the CD4 binding site. As observed for 3BNC117, disruption of Env-CD4 interaction in absence of Nef and Vpu (N-U-D368R) enhanced the recognition of infected CD4^+^ T cells by PGT151 but not in infected macrophages, consistent with low levels of CD4 in this cell type.

The bNAb 10E8 recognizes the membrane proximal external region (MPER) of gp41 (Huang et al., 2012). While binding the unliganded trimer likely explains its neutralization breath (Huang et al., 2012), 10E8’s epitope, as with other MPER Abs, is better exposed upon CD4 interaction (Dimitrov et al., 2007). Accordingly, Nef deletion enhanced recognition of CD4^+^T cells infected with the N- and N-U- constructs and was dependent on Env–CD4 interaction since introduction of the D368R mutation significantly decreased its binding (Figure 2H). Interestingly, no 10E8 recognition for macrophages was observed upon Nef and Vpu deletion (Figure 2H) despite enhanced recognition of CD4^+^T cells infected with the N- and N-U- constructs. For CD4^+^T cells this effect appeared to be dependent on Env–CD4 interaction since introduction of the D368R mutation significantly decreases its binding (Figure 2H). Whether this cell-type-dependent discrepancy is linked to the different levels of CD4 at the cell surface remains to be determined. Of note, however, similar phenotypes were observed with the three additional IMCs tested (JRFL, YU2 and CH77) (Figures S2-S4).

We then evaluated the recognition of infected cells by nnAbs. In agreement with the “closed” conformation adopted by the unliganded Env trimer, none of the tested nnAbs efficiently recognized WT infected cells (Figure 3A-D). The CD4-triggerable nature of the epitope they recognize has been well documented (Alsahafi et al., 2015; Ding et al., 2016a; Ding et al., 2016b; Gohain et al., 2016; Prévost et al., 2022; Prevost et al., 2018; Veillette et al., 2014). Accordingly, all Abs recognized more efficiently N-, U- and N- U- infected CD4^+^ T cells. Abrogation of Env-CD4 interaction by introduction of the D368R mutation significanly impaired recognition by these nnAbs (Figure 3), further supporting their CD4-induced nature. Consistent with lower CD4 levels on macrophages (Lee et al., 1999), these Abs barely detected infected macrophages with the exception of N- and N-U- infected cells.

**Figure 3.**
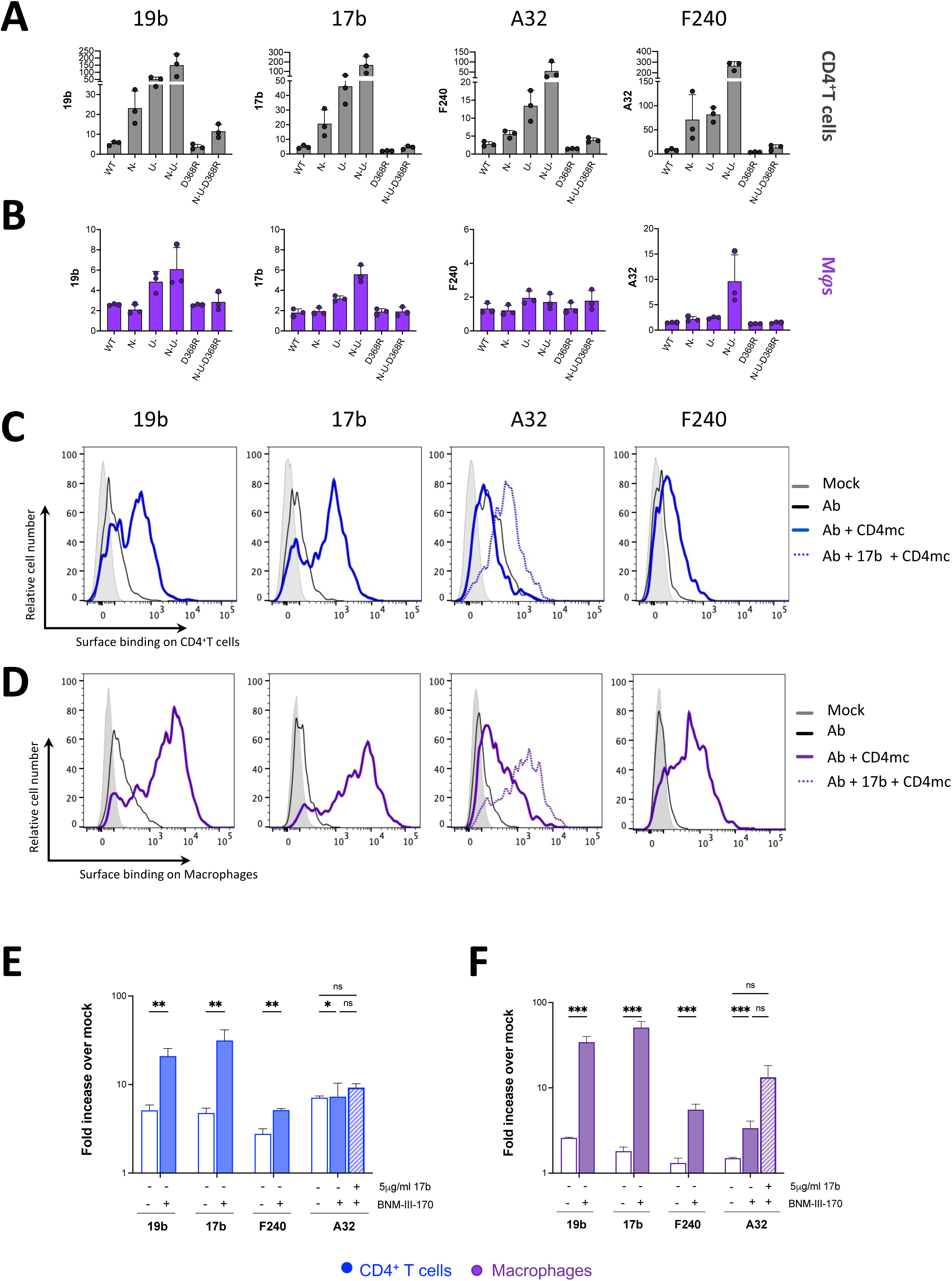
Recognition of HIV-1-infected primary CD4^+^ T cells and autologous macrophages by non-neutralizing antibodies. Autologous (**A** and **C**) CD4^+^T cells and (**B** and **D**) macrophages infected with the HIV- 1_AD8_ panel of viruses [wt, nef-defective (N-), Vpu-defective (U-), or both Nef and Vpu (N-U-) and CD4BS mutant (D368R) and N-U-D368R mutant] for 48h (CD4^+^ T cells) or 5 days (Macrophages) were stained with non-neutralizing antibodies 19b (V3 crown), 17b (Co-receptor binding site), A32 (Anti-cluster A) and F240 (gp41-Disulfide loop region). Histograms depict representative staining of infected cells of 19b, 17b, F240 and A32 with or without CD4mc BNMIII170 or A32 with 17b and CD4mc. Fold increase in nnAb binding for CD4^+^ T cells (**E**) and Macrophages (**F**) in the presence of the CD4mc is shown on the y-axis. Statistical significance was tested using Mixed Effect analysis (*p<0.05; **p<0.001; ***p<0.0001; ns, non-significant).

### Exposing CD4i epitopes on the surface of infected macrophages

It has been well documented that nnAbs fail to recognize Env in its “closed” conformation (Ding et al., 2016b; Prevost et al., 2017; Richard et al., 2016a; Richard et al., 2018; Richard et al., 2017; Richard et al., 2015; Richard et al., 2016b; Tolbert et al., 2016; Veillette et al., 2015; Veillette et al., 2014; von Bredow et al., 2016). As shown in Figure 3, infected macrophages make no exception with no recognition of wild-type infected cells by all tested nnAbs.

Small CD4-mimetic compounds (CD4mc) were shown to expose vulnerable Env epitopes at the surface of infected CD4^+^ T cells resulting in their sensitization to ADCC mediated by nnAbs and plasma from infected individuals (Alsahafi et al., 2019; Richard et al., 2016a; Richard et al., 2015). To evaluate whether this could apply to infected macrophages, we used BNM-III-170, a CD4mc extensively used to expose epitopes recognized by nnAbs on infected primary CD4^+^ T cells (Alsahafi et al., 2019; Anand et al., 2019; Madani et al., 2018; Rajashekar et al., 2021). To facilitate comparisons, we infected autologous primary CD4^+^T cells and evaluated the capacity of four non-neutralizing antibodies to recognize infected cells (macrophages and CD4^+^T cells). As described above, in the absence of CD4mc the four nnAbs tested (19b, 17b, A32 and F240) poorly recognized wild type infected macrophages or CD4^+^T cells (Figure 3A-B). Consistent with the capacity of BNM-III-170 to “open-up” Env and expose vulnerable epitopes, CD4mc addition enabled efficient recognition by 19b, 17b and F240 (Figure 3C-D). In both infected CD4^+^T cells and macrophages, and in agreement with the literature (Alsahafi et al., 2019; Anand et al., 2019; Rajashekar et al., 2021; Richard et al., 2016a), efficient exposure of the gp120 inner domain cluster A region recognized by A32, required addition of the CoRBS 17b antibody (Figure 3C-F, Figure S4). This was recapitulated with the other IMCs (Figure S4D-E).

### BNM-III-170 sensitizes HIV infected macrophages to ADCC mediated by HIV^+^ plasma

A recent study reported that infected macrophages are resistant to ADCC mediated by NK cells (Clayton et al., 2021). However, it remained to be determined whether CD4mc render infected macrophages susceptible to ADCC. To do this, we adapted our FACS-based ADCC assay that uses infected CD4^+^T cells (Richard et al., 2018; Veillette et al., 2014) for assessment of killing of infected macrophages (Figure 4, gating strategy shown in Figure S5). Briefly, macrophages differentiated for seven days were infected with HIV-1_AD8_ wild type and virus defective for Nef and Vpu (N-U-) for five days. Wild-type infected cells were then treated with or without BNM-III-170. N-U- infected cells were used as a positive control for ADCC activity mediated by nnAbs. ADCC activity was then measured following five hours incubation with autologous PBMCs as effector cells in the presence of plasma from nine different HIV-1 infected individuals Consistent with their role in protecting infected CD4^+^ T cells from ADCC, deletion of Nef and Vpu (N-U-) markedly increased their susceptibility to ADCC (Figure 4A). However, in infected macrophages deletion of Nef and Vpu only led to a small increase in ADCC susceptibility (Figure 4B). This is consistent with the modest increase in CD4 levels at the surface of N-U- compared to WT-infected macrophages (Figure 1A). Of note, addition of BNM-III-170 sensitized both infected CD4^+^T cells and macrophages to ADCC mediated by plasma from nine different HIV-1-infected individuals (Figure 4).

**Figure 4.**
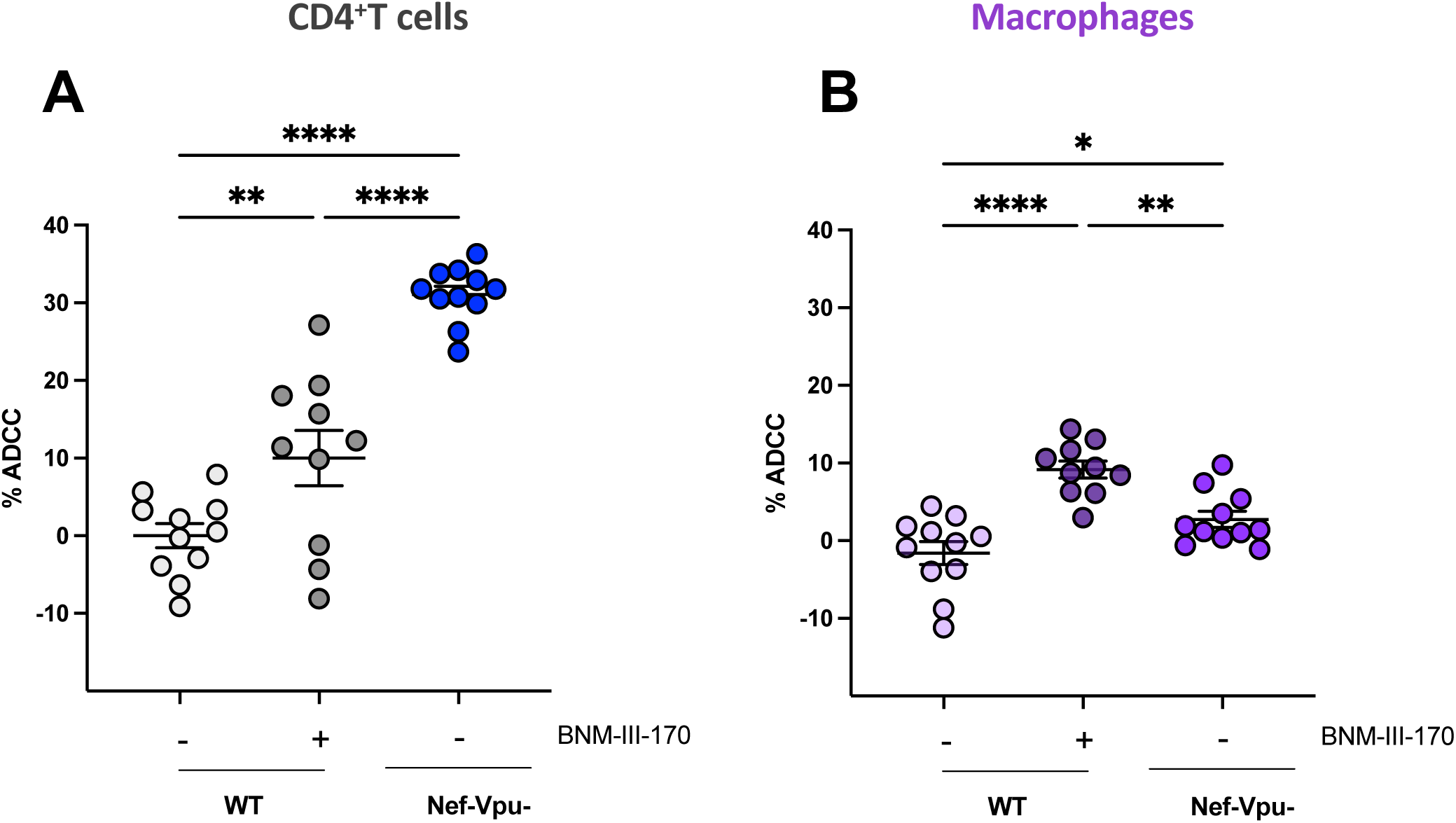
Small CD4mc BNM-III-170 sensitizes HIV-1-infected macrophages to ADCC mediated by HIV+ plasma. Autologous CD4^+^T cells and macrophages were infected with HIV-1_AD8_ [wt and nef- and Vpu-defective (N-U-) virus] and subsequently incubated with autologous PBMCs for 5h in the presence of HIV+ plasma with or without the CD4mc BNMIII170. Percent (%) ADCC of CD4^+^ T cells (**A**) and macrophages (**B**). Statistical significance was tested using Ordinary One-Way ANOVA followed by Holm-Šídáks multiple comparisons test (*p<0.05; **p<0.001; ***p<0.0001; ns, non-significant).

### Integrated analysis of the associations between recognition of Env epitopes, CD4 and BST-2 on the surface of macrophages and CD4^+^T cells

To assess the association of Env epitope recognition, CD4 and BST-2, we performed a network correlation analysis, done separately for CD4^+^T cells and macrophages (Figure 5). For macrophages, the network had two distinct clusters of significant correlations (Figure 5B), whereas the T cell network was more intertwined (Figure 5A). While there were significant associations between nnAb recognition in the presence of the CD4mc and the measurements of BST-2 and CD4, only two antibodies, A32 and F240, significantly correlated with BST-2 measurements (Figure 5B, *bottom panel*).

**Figure 5.**
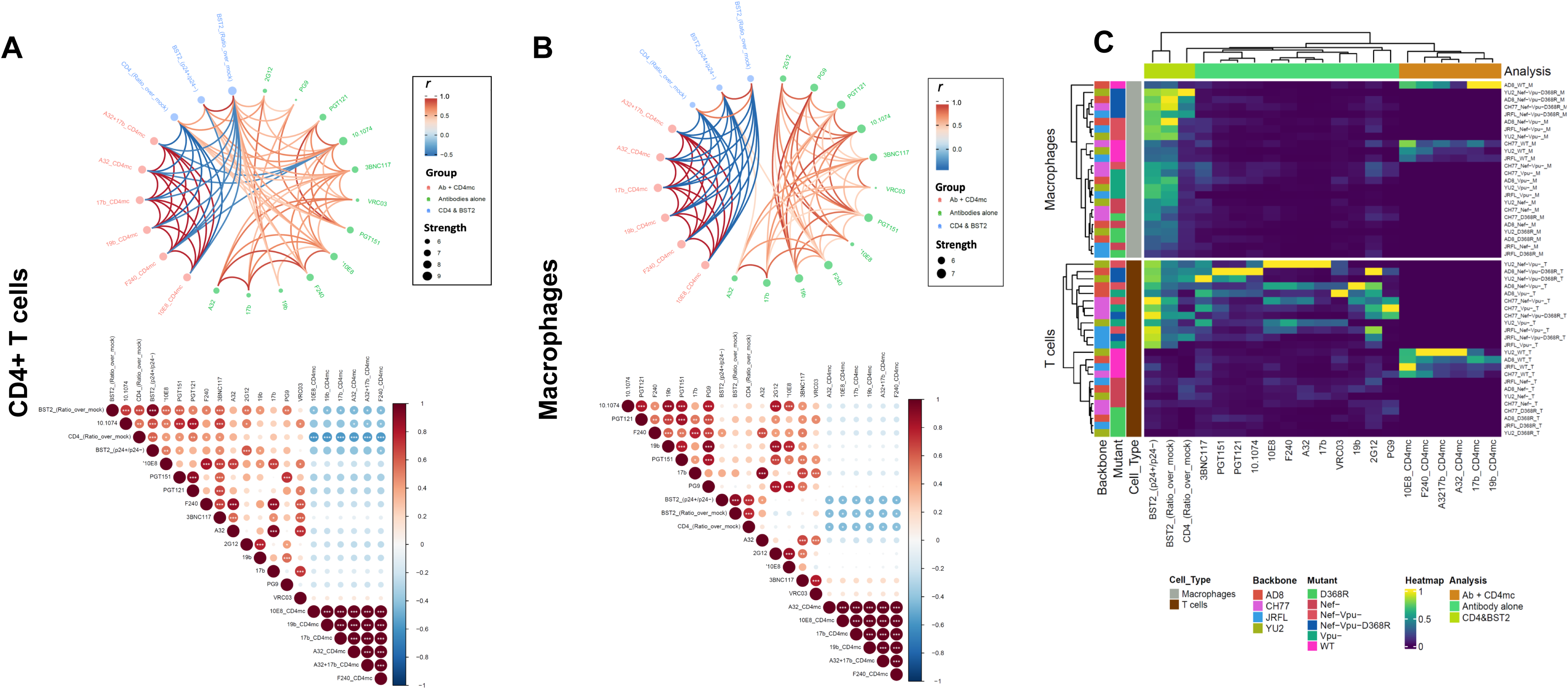
Network associations of Env epitope recognition with BST2 and CD4 on CD4^+^ T cells and macrophages. Circular edge bundling plots (top) and correlation matrices (bottom) for CD4^+^T cells (**A**) and Macrophages (**B**) where red and blue edges (top) and circles (bottom) represent positive and negative pairwise correlations between connected variables, respectively. Nodes in the edge bundling plots (top) are color-coded based on the grouping of variables shown in the legend with node size corresponding to degree of relatedness of correlations. Only significant correlations (p < 0.05, Spearman rank test) are displayed in the edge bundling plots (top). In the correlograms (bottom): * p < 0.05, ** p < 0.01, *** p < 0.005. (**C**) Heatmap showing Env, BST and CD4 expression relative to virus strains tested and cell type. Columns represent recognition of BST2, CD4 and Env clustered based on similarity and grouped by analysis method according to the color code provided. Rows represent the viruses/mutants grouped according to cell type. Virus IDs are clustered according to their binding profile, done separately for each cell type.

Importantly there was a striking disconnect of associations between bNAb binding and measurements of CD4 suggesting that Env epitope exposure on macrophages is minimally influenced by the surface levels of CD4. In stark contrast, the correlation network for CD4^+^T cells (Figure 5A, *top panel*) was more balanced with intricate relationships between the levels of CD4/BST-2 and Env recognition by all antibodies, including the nnAbs. Furthermore, the inverse correlations between the levels of CD4 and recognition by nnAbs in the presence of the CD4mc on CD4^+^T cells was more pronounced (Figure 5A *bottom panel*), supporting that the levels of CD4 on CD4^+^T cells greatly influence epitope recognition. Importantly, the heat map analysis (Figure 5C) highlights that the differences between macrophages and CD4^+^T cells are not virus strain dependent.

## Discussion

Recent work reported differential recognition of Env epitopes on the surface of infected macrophages (Clayton et al., 2021; Kek et al., 2021) with one study highlighting strong recognition of CD4BS epitopes on the surface of infected macrophages compared to CD4^+^T cells (Clayton et al., 2021). Here, we report that similarly to infected primary CD4^+^T cells, Env on macrophages is predominantly in a “closed” conformation.

HIV-1 uses three different proteins (Nef, Vpu and Env) to efficiently downregulate CD4 from the cell surface. While Nef targets CD4 molecules already present at the plasma membrane, Vpu and Env target newly-synthesized CD4 for degradation. While in both cell types deletion of Nef, Vpu and abrogation of Env-CD4 interaction (N-U- D368R) was required to restore CD4 levels to those observed with uninfected cells, the effect of Nef and Vpu appeared more prominent in CD4^+^T cells. Indeed, in this cell type but not in macrophages, their individual deletion (N- or U-) significantly increased CD4 levels. The relatively limited amount of CD4 at the surface of macrophages compared to autologous CD4^+^ T cells (Figure 1A and (Lee et al., 1999)) might facilitate the task of the two other remaining viral proteins to fully downregulate CD4 in macrophages.

In agreement with previous studies (Prévost et al., 2022; Prevost et al., 2018; Veillette et al., 2014), we found that the ability of Nef and Vpu to protect infected cells from ADCC was linked to CD4 expression. Both CD4^+^ T cells and macrophages infected with the wild-type virus were resistant to ADCC. However, while Nef and Vpu deletion dramatically increased the susceptibility of infected CD4^+^ T cells to ADCC, it only had a minor effect in infected macrophages (Figure 4). This is consistent with the limited amount of CD4 present at the surface of infected macrophages to “open-up” Env and expose ADCC vulnerable epitopes. Nevertheless, similarly to infected primary CD4^+^ T cells, we were able to expose vulnerable epitopes at the surface of infected macrophages using the CD4mc BNM-III-170. Of note, exposure of the gp120 inner domain A32 epitope further required the combination of the CD4mc with a CoRBS Ab such as 17b. This combination was recently shown to stabilize an ADCC vulnerable conformation of Env, State 2A (Alsahafi et al., 2019), indicating that this conformation can also be stabilized on infected macrophages. Interestingly, the combination of nnAbs (17b + A32) and CD4mc was also recently found to decrease HIV-1 replication and reduce HIV DNA in CD4^+^T cells in humanized mice (Rajashekar et al., 2021). Accordingly, the data herein demonstrate that the addition of BNM-III-170 improved recognition and ADCC susceptibility of HIV-1 infected macrophages by nnAbs, commonly elicited antibodies that predominate in HIV+ plasma. These findings warrant further efforts to test whether CD4mc could enable nnAbs to eliminate infected macrophages *in vivo*.

Since a recent study reported that macrophages are resistant to NK cell killing (Clayton et al., 2021), we decided to perform our ADCC assay using PBMCs as effector cells. Our results suggest that infected macrophages could be sensitive to Fc-effector functions mediated by other immune cells present in peripheral blood. Here, we used MDM as a model, it would therefore be important to ascertain whether the phenotypes observed herein are consistent with other cells of the myeloid lineage, in particular microglia, an important HIV reservoir within the central nervous system (CNS) (Wallet et al., 2019). While we show that the small molecule CD4mc BNM-III-170 was able to expose CD4i epitopes on infected macrophages permitting ADCC, we acknowledge that the CNS may still remain inaccessible.

Altogether, our data provides new insights on the role of viral proteins in modulating cell-surface CD4 and Env conformation on infected macrophages, as well as proof of principle that HIV-1-infected macrophages can be sensitized to recognition and ADCC by CD4i antibodies. These findings provide new information for the development of immunotherapies aimed at targeting and eliminating the HIV-1 reservoir *in vivo*.

### Limitations of the study

While we are able to show that HIV-1 infected macrophages can be made susceptible to ADCC mediated killing, it is important to establish the effector cells within the PBMC pool responsible for this phenotype. Future work will therefore be undertaken to identify the cell type(s) responsible for the elimination of infected macrophages. Additionally, here we used MDM as a model and it would be important to extend our findings to additional macrophage subtypes, from different lineages. Nevertheless, the data presented here using MDMs facilitates important assertions as to the mechanism of macrophage evasion of immune responses.

## Acknowledgment

The authors thank the CRCHUM BSL3 and Flow Cytometry Platforms for technical assistance, Mario Legault from the FRQS AIDS and Infectious Diseases network for cohort coordination and clinical samples. We thank Dennis Burton (The Scripps Research Institute) for the JR-FL infectious molecular clone and Michel Nussenzweig for 3BNC117 and 101074 antibodies. This study was supported by grants from the National Institutes of Health to A.F., and J.C.K. (R01 AI148379), to A.F. (R01 AI129769 and R01 AI150322), to B.H.H. (R01 AI162646 and UM1 AI164570) and the Basic Science Core of the University of Alabama at Birmingham Center for AIDS Research (AI27767). This work was also partially supported by 1UM1AI164562-01, co-funded by National Heart, Lung and Blood Institute, National Institute of Diabetes and Digestive and Kidney Diseases, National Institute of Neurological Disorders and Stroke, National Institute on Drug Abuse and the National Institute of Allergy and Infectious Diseases, a CIHR foundation grant #352417, a CIHR Team grant #422148 and a Canada Foundation for Innovation grant #41027 to A.F. A.F. is the recipient of a Canada Research Chair on Retroviral Entry #RCHS0235 950-232424. F.K. is supported by the Deutsche Forschungsgemeinschaft (CRC 1279 and SPP 1923). A.L. and R.G. were supported by a MITACS Accélération postdoctoral fellowship. The funders had no role in study design, data collection and analysis, decision to publish, or preparation of the manuscript.

## Author contributions

A.L. and A.F. conceived the study. A.L., S.D. and A.F. designed experimental approaches. A.L., L.M., S.D, H.M., D.C., G.B.B., R.G., G.G.L., J.R., R.D., and A.F. performed, analyzed, and interpreted the experiments. B.H., F.K., J.C.K., H.D., H.C.C., A.B.S. supplied novel/unique reagents. A.L, B.H.H., and A.F. wrote the paper. All authors have read, edited, and approved the final manuscript.

## Declaration of Interests

The authors declare no competing interests

## STAR METHODS

### RESOURCE AVAILABILITY

#### Lead contact

Further information and requests for resources and reagents should be directed to and will be fulfilled by the lead contact, Andrés Finzi.

#### Materials Availability

All other unique reagents generated in this study are available from Andrés Finzi with a completed Materials Transfer Agreement.

#### Data and code availability

- All data reported in this paper will be shared by the lead contact upon request.
- This paper does not report original code.
- Any additional information required to reanalyze the data reported in this paper is available from the lead contact upon request.

### EXPERIMENTAL MODEL AND SUBJECT DETAILS

#### Ethics Statement

Written informed consent was obtained from all study participants and research adhered to the ethical guidelines of CRCHUM and was reviewed and approved by the CRCHUM Institutional Review Board (ethics committee, approval number CE16.164- CA). Research adhered to the standards indicated by the Declaration of Helsinki. All participants were adult and provided informed written consent prior to enrolment in accordance with Institutional Review Board approval.

#### Cells

The HEK293T human embryonic kidney cells (293T) (obtained from ATCC and the NIH AIDS Research and Reference Reagent Program, respectively) were maintained at 37°C and 5% CO_2_ in Dulbecco’s modified Eagle’s medium (DMEM) (Invitrogen) supplemented with 5% FBS (VWR) and 100mg/ml of penicillin-streptomycin (Wisent). Primary cells were grown as previously described (Veillette et al., 2014). Briefly cryopreserved human peripheral blood mononuclear cells (PBMCs) isolated by ficoll density gradient from 6 healthy donors (HIV and hepatitis C virus [HCV] seronegative) (4 males and 2 females), who gave written informed consent under research protocols approved by the CRCHUM, were thawed and monocytes were isolated by plate adherence in 10cm petri dishes (Sarsdedt) for 1h in Iscove’s modified Dulbecco medium (IMDM). Non-adherent cells were collected while adherent cells washed extensively in serum free media and allowed to differentiate to macrophages for seven days in IMDM supplemented with 100mg/ml of penicillin-streptomycin and 10% pooled human sera (Valley Biomedicals), with a half media change at day 3. CD4^+^T cells were isolated from the non-adherent cells by negative selection (EasySep) activated and maintained in culture as previously described (Prévost et al., 2020; Richard et al., 2016b).

### METHOD DETAILS

#### Infectious molecular clones (IMCs) and plasmids

The infectious molecular clones (IMCs) of HIV-1 strain AD8: (pAD8^+^) HIV-1AD8 (Theodore et al., 1996) Vpu- (AD8- U-), Nef- (AD8-N-) and Nef-Vpu- (AD8-N-U-); and HIV-1 YU2 strain: HIV-1 YU2 (Li et al., 1991), Vpu- (YU2-U-) and Nef-Vpu- (AD8-N-U-). The D368R derivatives of AD8 and YU2 and Nef defective YU2 (YU2-N-) were generated by mutagenesis using primers listed in Key resource table. The IMCs of HIV-1 strains JRFL was kindly provided by Dr Dennis Burton. The Nef-defective and Vpu-defective JR-FL IMCs were previously described (Prévost et al., 2022). The CH77 (CH077) transmitter founder was previously described (Ochsenbauer et al., 2012). Mutated IMCs were previously described (Bar et al., 2012; Fenton-May et al., 2013; Heigele et al., 2016; Ochsenbauer et al., 2012; Richard et al., 2015).

#### Antibodies and HIV+ plasma

The following antibodies were used to assess cell surface staining: 2G12, PG9, PGT121, 10.1074, 3BNC117, VRC03, PGT151, 10e8, 19b, 17b, A32, F240. Details of which are listed in the key resources table. The panel of anti-HIV antibodies were conjugated with CF647 probe (Sigma Aldrich) as per the manufacturer instructions and used for cell-surface staining of HIV-1-infected primary CD4^+^T cells and macrophages. Mouse anti-human CD4 (Clone OKT4, BV421- conjugated; Biolegend, San Diego, CA, USA) and mouse anti-human BST2 (clone RS38E, PE-Cy7-conjugated; BioLegend, San Diego, CA, USA) were used as primary antibodies for cell surface staining. To confirm purity of macrophages the mouse anti-human CD3 (Clone UCHT1; BUV395 conjugated; BD Biosciences) and to confirm differentiation of monocyte differentiation to macrophages the Rat anti-human/mouse CD11b (BV650 conjugated; Biolegend) were also used. Plasma used for ADCC experiments were collected from HIV-infected, heat-inactivated for 1h at 56°C and conserved as previously described (Prévost et al., 2022) until ready to use in subsequent experiments.

#### CD4-mimetic BNM-III-170

The small molecule CD4-mimetic BNM-III-170 synthesized as previously described (Melillo et al., 2016) was dissolved in dimethyl sulfoxide (DMSO) at a stock concentration of 10 mM, aliquoted, and stored at − 20 °C prior to being diluted to 50μM in PBS for cell-surface staining and ADCC assays.

#### Viral stock production, Infections and Detection of Infected Cells

Vesicular stomatitis virus G (VSVG)-pseudotyped viruses were produced in 293T by Polyethylenimine (PEI) transfection. Briefly 1 x10^7^ 293T seeded in a TC75cm^2^ flasks (Cat# 734-2315; VWR) were transfected with plasmids encoding VSVG and HIV infectious molecular clone (IMC) at a ratio of 2:5. 4h post transfection, the supernatant was removed and replaced with fresh media and cultured for a further 72h. Cell supernatants were harvested, clarified by centrifugation, and filtered through a 0.45µM filter (Minisart) and layered onto a 25% sucrose gradient prior to ultracentrifugation. Pseudotyped viruses (concentrated 100-fold) were subsequently titrated on CD4^+^T cells as described (Veillette et al., 2015). In addition day 7 differentiated macrophages were cultured in 3.5cm petri dishes and passively infected for 5 days to determine macrophage specific inoculum required to obtain >10% infection.

#### Flow Cytometry Analysis of Cell-surface Staining and ADCC Responses

Five days post-infection, macrophages were washed in PBS, incubated in 10 mM EDTA for 30 minutes at RT, detached and transferred to 96-well V-bottom plates (Corning; Cat # 0877126). In parallel, CD4^+^T cells infected for 48h were harvested, washed and transferred to 96-well V-bottom plates and incubated for 30 minutes with AquaVivid viability dye (Thermo Fisher Scientific, Cat# L43957) as per manufacturer’s instructions. Cells were then washed twice in PBS. Prior to staining with antibodies, macrophages were incubated with 10% human sera (Valley Biomedicals) and 2% FcBlock (Miltenyi) in FACS buffer (1% BSA, 1mM EDTA in PBS) for 10 minutes. Following Fc blocking, macrophages and CD4^+^T cells resuspended in 1% BSA were incubated with a panel of anti-Env antibodies (Table S2; Key Resources table) pre-coupled to CF647 fluorophore (Sigma-Aldrich) for 30 minutes at RT. Binding of 19b, 17b, A32, F240 and 10e8 was done with or without BNM-III-170 (50μM). Cells were then washed twice with FACS buffer, fixed with 2% Paraformaldehyde (PFA) and permeabilized using BD CytoFix/CytoPerm Fixation/Permeabilization Kit (BD Biosciences) as per manufacturer’s instructions. Detection of p24+ infected cells was performed as described (Richard et al., 2016b; Veillette et al., 2015) using anti-Gag p24-FITC (Beckman Coulter). The percentage of infected cells (p24 + cells) was determined by gating the living cell population based on the viability dye staining (Aqua Vivid, Thermo Fisher Scientific, Cat# L43957). Samples were analyzed on a Fortessa cytometer (BD Biosciences, Mississauga, ON, Canada) and data analysis was performed using Flow Jo version 10.4.0 (Tree Star, Ashland, OR, USA).

Measurement of ADCC-mediated killing was performed as previously described (Prévost et al., 2022; Richard et al., 2015). Briefly, primary CD4^+^ T cells infected for 48h with the AD8 wild type (WT) virus of viruses defective for Nef and Vpu (N-U-) were co-cultured with autologous PBMC (Effector: Target ratio of 10:1) in the presence of HIV+ plasma (1:1000) with or without BNM-III-170. Macrophage ADCC assays were performed as above with the following modifications; macrophages infected for 5 days with AD8 WT or N-U- were incubated with 10% human sera prior to co-culture with PBMCs and HIV+ plasma (1:1000) with or without BNM-III-170 for 5h at 37°C. The percentage of cytotoxicity was calculated as described (Prévost et al., 2022; Richard et al., 2015).

#### Statistical Analyses and Visualization

Statistics were analyzed using GraphPad Prism version 9.3.1 (GraphPad, San Diego, CA, USA). All data sets with statistical analysis was tested for statistical normality and this information was used to apply the appropriate statistical tests. P values < 0.05 were considered significant; significance values are indicated as *p < 0.05, **p < 0.01, ***p < 0.001, ****p < 0.0001. **C**ircular edge bundling graphs were generated in undirected mode in program R v.4.1.2 (R Core Team., 2017) using ggraph, igraph, tidyverse, and RColorBrewer packages. Edges are only shown if p < 0.05, and nodes are sized according to the sum of the connecting edges’ absolute r values. Nodes are color-coded according to the groups of variables. Correlograms were generated using the corrplot and RColorBrewer packages using hierarchical clustering based on the first principal component. Normalized heatmaps with dendrograms were created using the complexheatmap and tidyverse packages. Normalizations were done per column (analysis).

## SUPPLEMENTAL INFORMATION

Supplemental information can be found online at …

---

**Figure S1.**
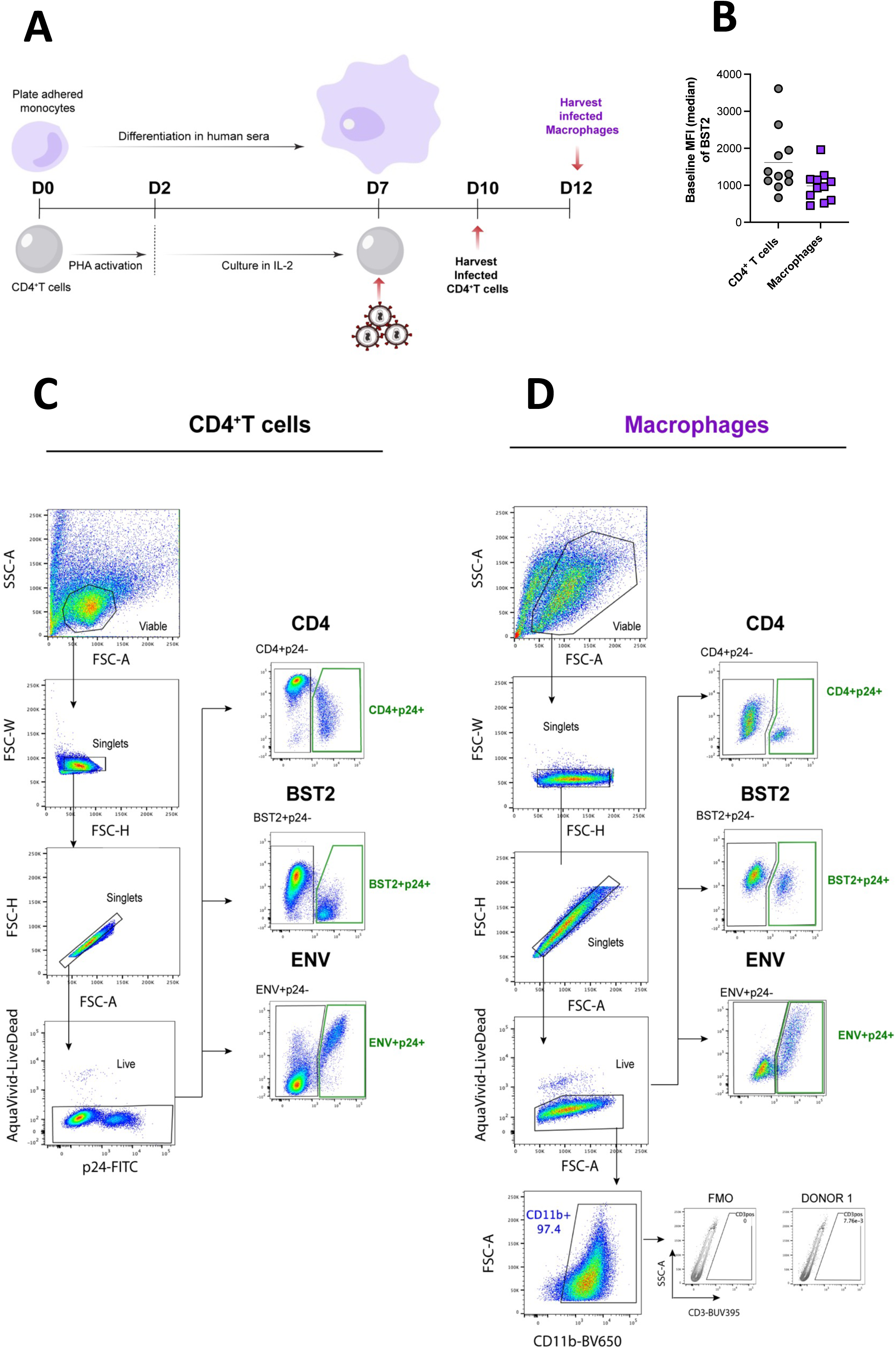
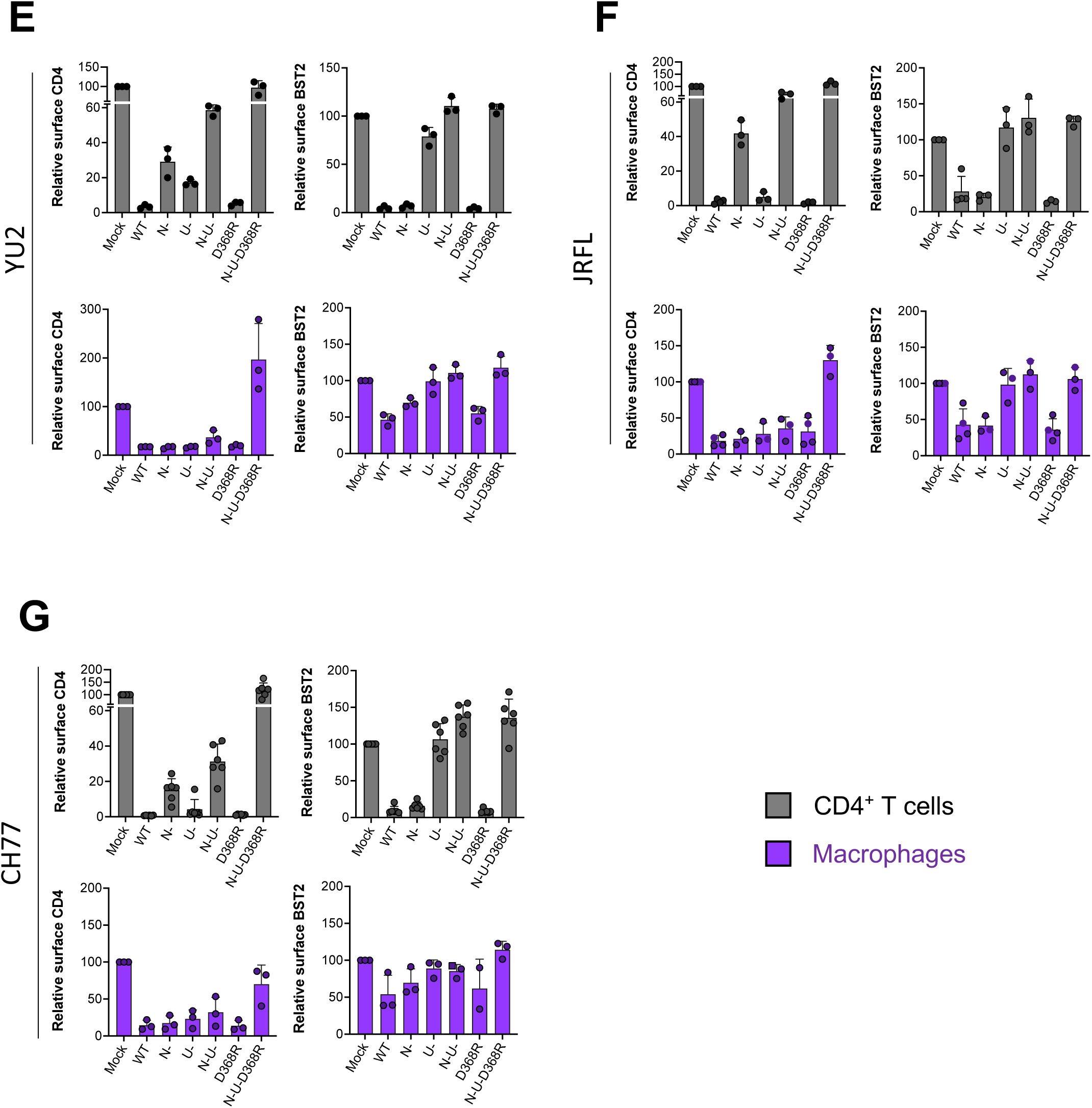
Effect of Nef, Vpu and Env on the surface expression of CD4 and BST2 in autologous CD4^+^ T cells and macrophages infected with different HIV-1 strains (Related to Figure 1). (*A*) Schematic of experimental timeline. Details are described in STAR METHODS. (*B*) Baseline Mean Fluorescence Intensity (MFI) of BST2 on uninfected CD4^+^ T cells and macrophages from 12 different experiments. Gating strategy for measurement of expression of Env, BST and CD4 on (*C*) CD4^+^ T cells and (*D*) macrophages; (*E-G*) Relative surface levels of CD4 and BST2 on autologous CD4^+^T cells and macrophages infected with panel of viruses [wt, nef-defective (N-), Vpu-defective (U-), or both Nef and Vpu (N-U-), CD4BS mutant (D368R) and N-U-D368R mutant] from (*E*) HIV-1_YU2_ (*F*) HIV-1_JR-FL_ and (*G* )HIV-1_CH77_

**Figure S2.**
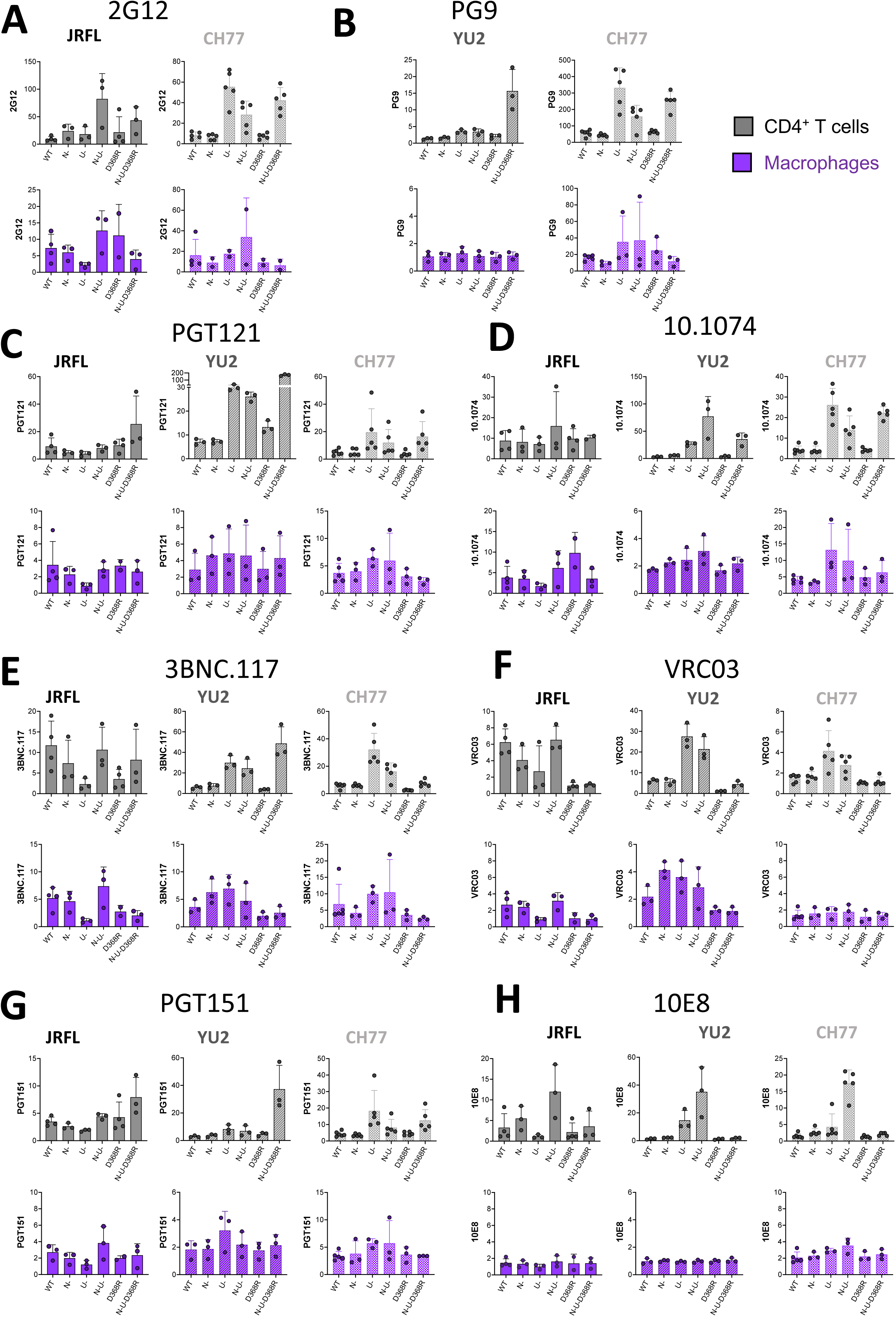
HIV-1 Env expression on the surface of autologous CD4^+^ T cells and macrophages infected with different HIV-1 strains. (Related to Figure 2). Autologous CD4^+^T cells and macrophages were infected with panel of viruses [wt, nef-defective (N-), Vpu-defective (U-), or both Nef and Vpu (N-U-) and CD4BS mutant (D368R) and N-U-D368R mutant] from HIV-1_JR-FL_, HIV-1_CH77_ and HIV-1_YU2_. 48h later (CD4^+^ T cells) or 5 days post-infection (Macrophages). Cells were then stained with (*A*) 2G12 a conformation independent antibody; (*B*) the V1V2-apex antibody PG9; V3glycan antibodies (*C*) PGT121 and (*D*) 10.1074; CD4BS antibodies (*E*) 3BNC117 and (*F*) VRC03; *(G)*PGT151 which targets the gp120-gp41 interface; and the *(H)* anti-MPER antibody 10E8.

**Figure S3.**
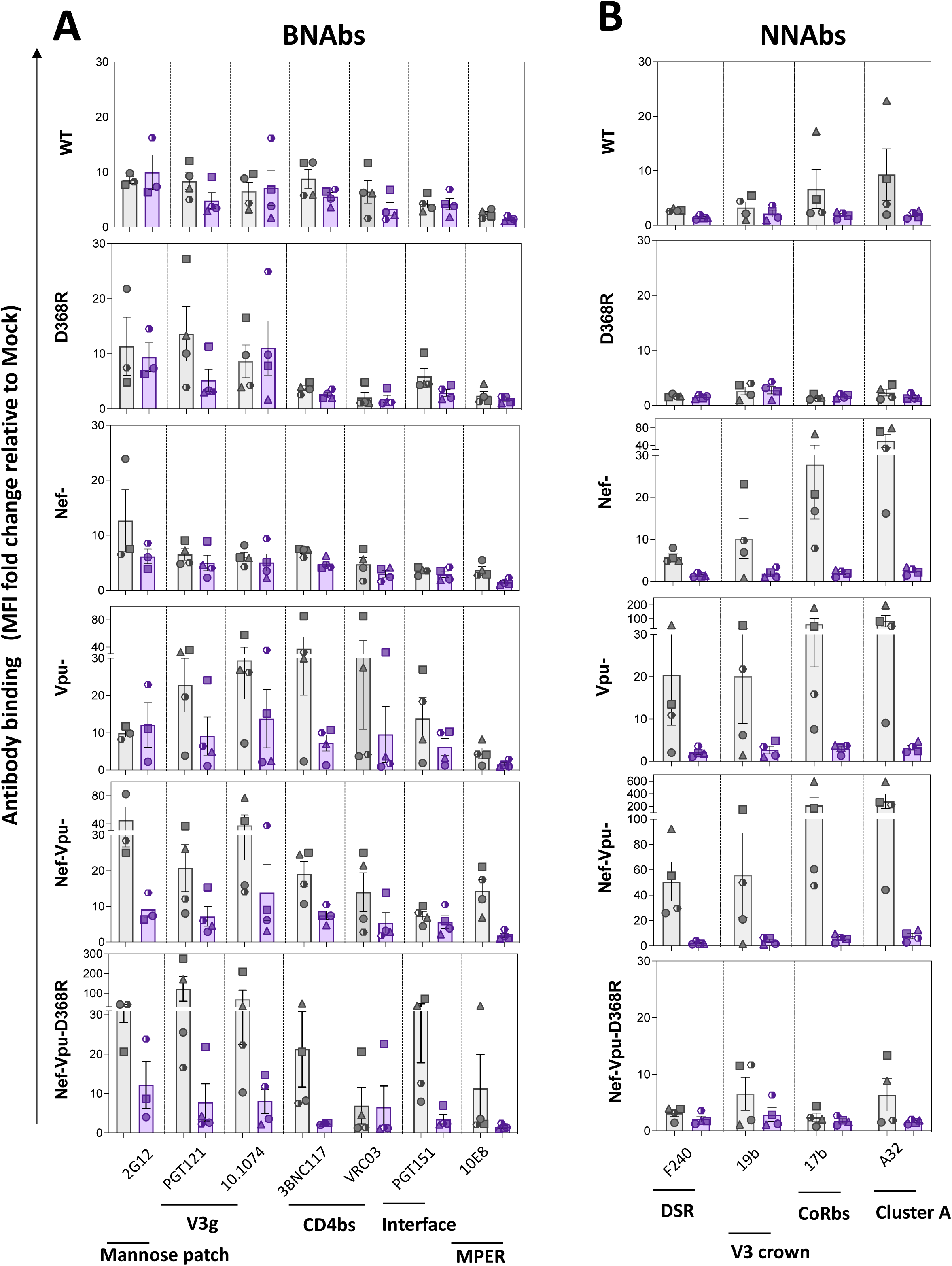
HIV Env conformational landscape on the surface of autologous CD4^+^ T cells and macrophages infected with different HIV strains (Related to Figures 2 and 3) Summary of Env recognition on Autologous (CD4^+^T cells and macrophages infected for 48h (CD4^+^ T cells) or 5 days (Macrophages) with panel of viruses [wt, nef-defective (N-), Vpu-defective (U-), or both Nef and Vpu (N-U-) and CD4BS mutant (D368R) and N-U-D368R mutant] of different IMCs (AD8, YU2, JRFL, CH77). *(A)* Recognition by bNAbs. *(B)* Recognition by nnAbs. Statistical significance was tested using Mixed Effect analysis (*p<0.05; **p<0.001; ***p<0.0001; ns, non-significant).

**Figure S4.**
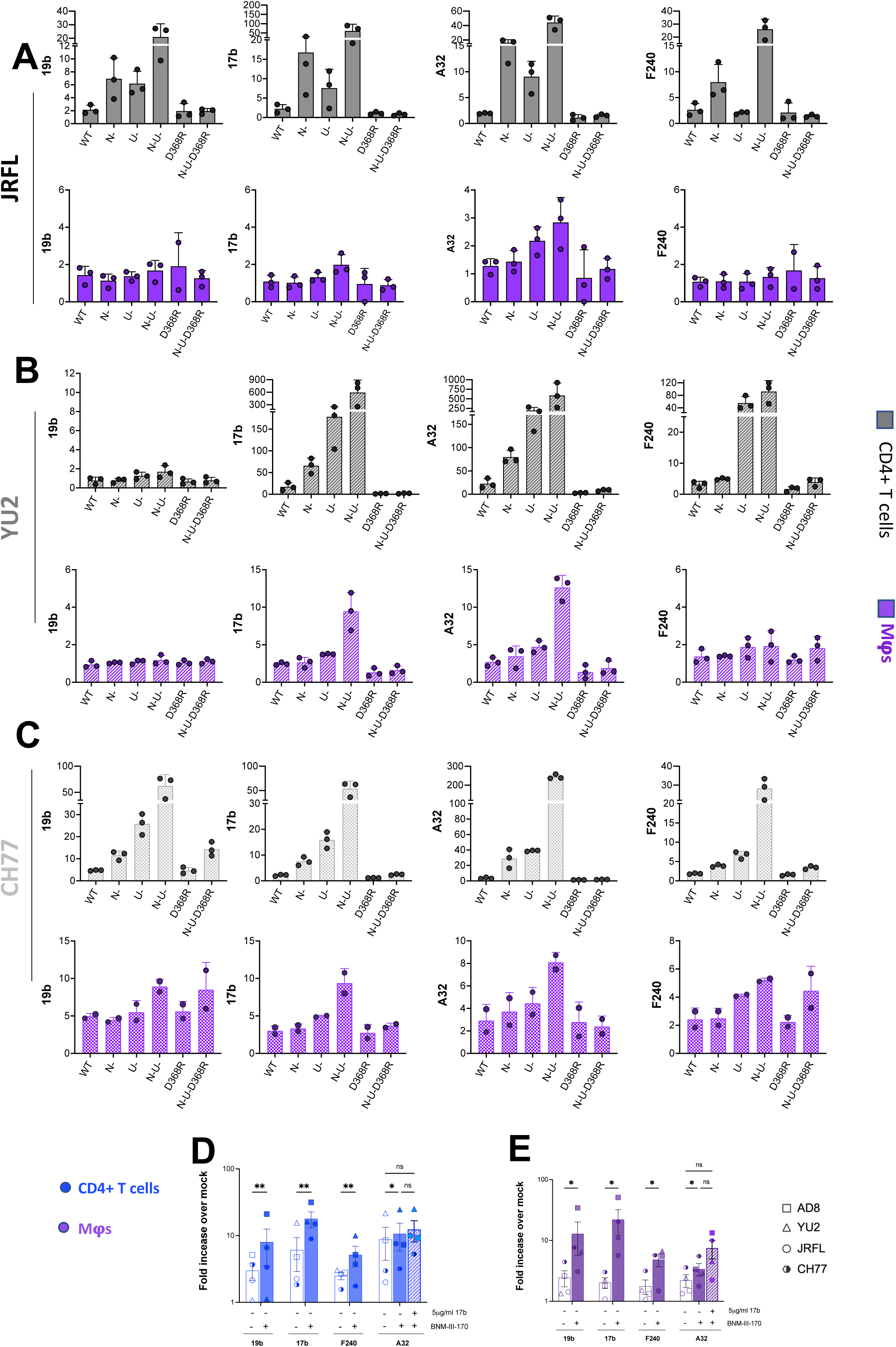
Small molecule CD4mc, BNM-III-170 opens Env to reveal epitopes recognized by nnAbs in CD4^+^T cells and macrophages infected with different HIV strains (Related to Figure 3). Autologous CD4^+^T cells and macrophages were infected with panel of viruses [wt, nef-defective (N-), Vpu-defective (U-), or both Nef and Vpu (N-U-) and CD4BS mutant (D368R) and N-U- D368R mutant] from (*A*) HIV-1_JR-FL_, (*B*) HIV-1_YU2_ and (*C*) HIV-1_CH77_. 48h later (CD4^+^ T cells) or 5 days post-infection (Macrophages), cells were stained with non-neutralizing antibodies 19b (V3 crown), 17b (Co-receptor binding site), A32 (Anti-cluster A) and F240 (gp41-Disulfide loop region). Fold increase in nnAb binding in the presence of the CD4mc for (*D*) CD4^+^T cells and *(E)* macrophages.

**Figure S5.**
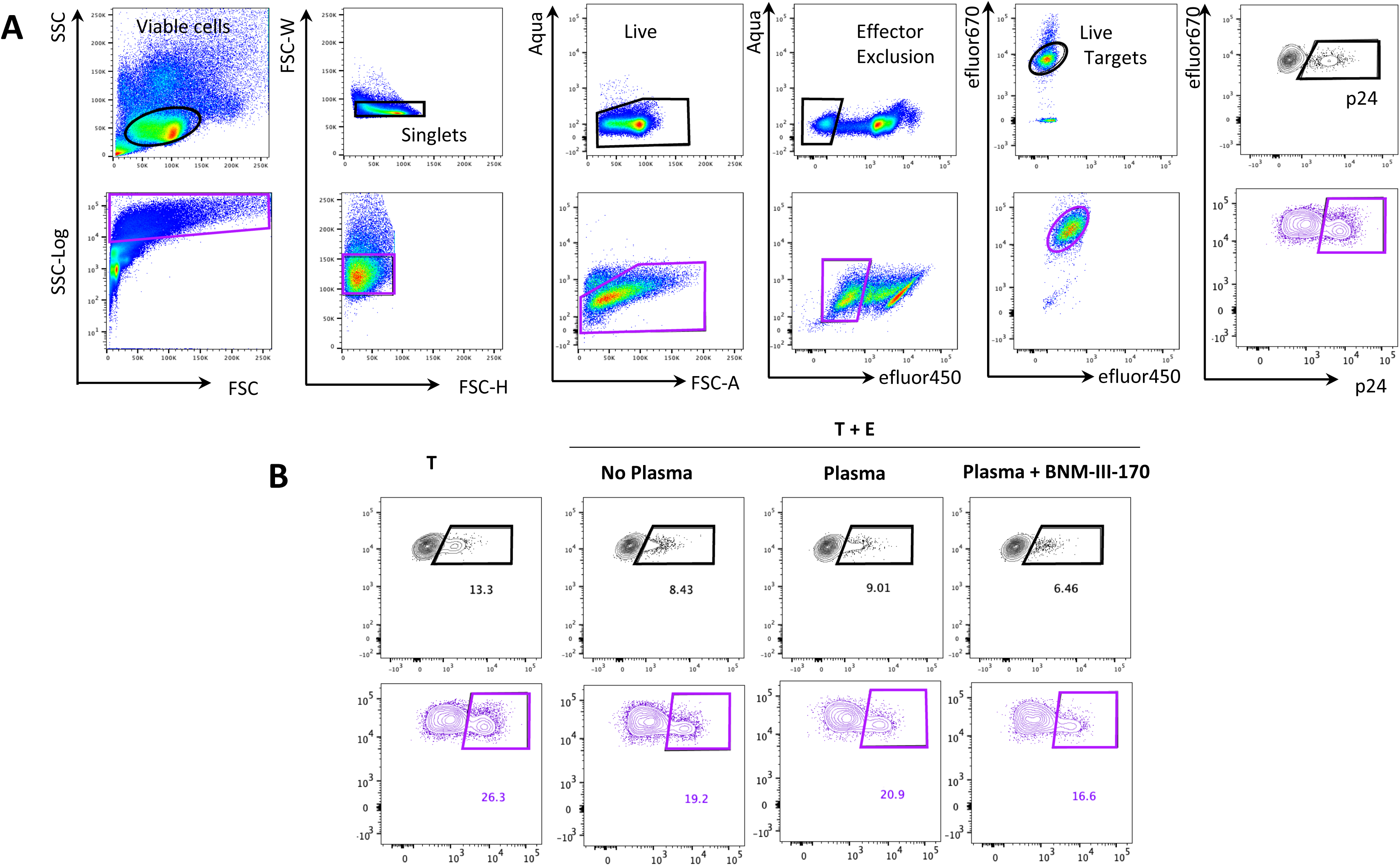
Gating strategy for CD4^+^T cells and Macrophages ADCC Assays (Related to Figure 4). Gating strategy for ADCC assays performed for CD4^+^T cells (*Top panel*) and Macrophages (*Bottom panel*) infected with HIV-1_AD8_ wildtype (WT) or virus defective to Nef and Vpu (N-U-) and incubated with autologous PBMCs for 5h. Macrophages and CD4^+^T cells were gated for viable cells. Side Scatter Axis for macrophages were transformed to log axis to visualize both the large macrophage targets and the small PBMC effectors. Subsequent doublet exclusion was performed, followed by live dead exclusion and effector exclusion. Targets were then selected and subsequent gating was done on the p24+ population. *(B)* Representative plots for the p24+ gate for Targets alone (T), and Targets with Effectors (T+E) in presence or absence of HIV+ plasma and/or CD4mc BNM-III-170

## Notes

### Competing Interest Statement

The authors have declared no competing interest.

